# Transfer of Knowledge from Model Organisms to Evolutionarily Distant Non-Model Organisms: The Coral *Pocillopora damicornis* Membrane Signaling Receptome

**DOI:** 10.1101/2021.10.18.464760

**Authors:** Lokender Kumar, Nathanael Brenner, Sam Sledzieski, Monsurat Olaosebikan, Matthew Lynn-Goin, Hollie Putnam, JK Yang, Nastassja Lewinski, Rohit Singh, Noah M. Daniels, Lenore Cowen, Judith Klein-Seetharaman

## Abstract

With the ease of gene sequencing and the technology available to study and manipulate non-model organisms, the need to translate our understanding of model organisms to non-model organisms has become an urgent problem. For example, mining of large coral and their symbiont sequence data is a challenge, but also provides an opportunity for understanding functionality and evolution of these and other non-model organisms. Much more information than for any other eukaryotic species is available for humans, especially related to signal transduction and diseases. However, the coral cnidarian host and human have diverged over 700 million years ago and homologies between proteins are therefore often in the gray zone or undetectable with traditional BLAST searches. We introduce a two-stage approach to identifying putative coral homologues of human proteins. First, through remote homology detection using Hidden Markov Models, we identify candidate human homologues in the cnidarian genome. However, for many proteins, the human genome alone contains multiple family members with similar or even more divergence in sequence. In the second stage, therefore, we filter the remote homology results based on the functional and structural plausibility of each coral candidate, shortlisting the coral proteins likely to be true human homologues. We demonstrate our approach with a pipeline for mapping membrane receptors in humans to membrane receptors in corals, with specific focus on the stony coral, *P. damicornis*. More than 1000 human membrane receptors mapped to 335 coral receptors, including 151 G protein coupled receptors (GPCRs). To validate specific sub-families, we chose opsin proteins, representative GPCRs that confer light sensitivity, and Toll-like receptors, representative non-GPCRs, which function in the immune response, and their ability to communicate with microorganisms. Through detailed structure-function analysis of their ligand-binding pockets and downstream signaling cascades, we selected those candidate remote homologues likely to carry out related functions in the corals. This pipeline may prove generally useful for other non-model organisms, such as to support the growing field of synthetic biology.

## INTRODUCTION

A bioinformatics functional genomics/proteomics pipeline for a newly sequenced non-model eukaryotic organism can (and should) seek to leverage the wealth of known information and annotation available for model species that are evolutionarily related. Due to the urgency of tackling species loss through environmental damage, many new non-model organisms are currently being sequenced, especially corals. Once the likely constituent genes have been identified (Ekblom & Wolf, 2014)(Campbell et al., 2014), predicting if the relevant evolutionary functions of the non-model species genes are conserved or have diverged from their orthologous counterparts in the model species becomes directly relevant. Of course, there may be genes in the non-model organism that have no homologues, also referred to as the dark genome (Oprea, 2019). However, the goal of this paper is to maximally exploit existing knowledge from model organisms for the mining of non-model organisms for function.This is because this step is often quite difficult for organisms such as corals, where the host is 700 million years distant from the closest well-annotated model organism, even for many of the genes in closely related species to the model organism (Copper, 2001). It is particularly interesting that coral animals contain a surprising number of human homologs missing from fly and *C. elegans* (Kortschak et al., 2003).

Corals are complex organisms consisting of an animal host (cnidarian) and a microbiome with more than 20,000 species consisting of symbiotic algae, as well as bacteria, bacteriophages, fungi and viruses, collectively referred to as holobiont (Thompson et al., 2014). We consider here, as a case study, the membrane receptor proteins in corals. These are important families of proteins to study in corals because they have large potential to mediate the coral animal’s interactions with its symbiont, microbiome and environment. Understanding these mechanisms in corals has become urgent. As a result of anthropogenic activities, both local and global, coral holobionts are declining rapidly. Mass coral bleaching, or the expulsion of the symbiotic algae due primarily to thermal stress driven by marine heatwaves, is resulting in substantial coral mortality (Hughes et al., 2018). A recent study assessed 100 worldwide locations and found that the annual risk of coral bleaching has increased from an expected 8% of locations in the early 1980s to 31% in 2016 (Hughes et al., 2018). Thus understanding the relevant parts of the coral animal genome that are relevant to interactions with their environment has become an urgent bioinformatics task, made possible by the recent availability of coral genomes for multiple species of both animal and symbiont. Fortunately, large classes of membrane receptor proteins have been preserved over time (de Mendoza et al., 2014), so there is a large body of prior knowledge that we can extrapolate from.

Given a protein of interest that has not been studied before, how much can be learnt from better studied proteins is a common question in biology. The answer lies in the sequence-structure-function paradigm, namely that a similar sequence usually maps to a similar structure and conserved function. This is the premise for the use of sequence alignments, for which many methods of analysis exist, the choice of which typically depends on how similar the sequences to be compared are. The problem with the sequence-structure-function paradigm is that relatively small changes in sequence can have dramatic effects on function, which is the basis for the evolution of protein families, where the overall fold is highly conserved, yet different members of the family can have different functions.

For many years, BLAST (Altschul et al., 1997) was the *de facto* standard homology search tool, as it provided comparable sensitivity to exact alignment methods such as Smith-Waterman (Smith & Waterman, 1981) and early heuristics such as FASTA (Pearson, 1991) with much faster runtime performance. However, these methods, and even iterative methods such as PSI-BLAST, are unable to detect homologs in the “twilight zone” of homology, between 10 and 30% sequence identity (Rost, 1999; Yona & Levitt, 2002). The development of profile hidden Markov models (HMMs) such as HMMER (Eddy, 1998) and SAM (Karplus et al., 1998) improved remote homology detection performance, but as they still attempt to score a single query sequence against a profile-based HMM, they may miss homologs deep into the “twilight zone.” A more recent advancement came in the form of HMM-HMM alignment (Söding et al., 2005) with HHpred in 2005, with further improvements resulting in HHblits (Remmert et al., 2011). These methods build a library of HMMs (or use an existing library) from families of nucleotide or protein sequences, and next build a *query* HMM from a multiple sequence alignment based on a BLAST-like search for similar sequences to the query, and use a variant of the Viterbi algorithm (Viterbi, 2009) to align the two HMMs according to maximum likelihood. HHblits was our choice for remote homology detection in this pipeline, as it combines state-of-the-art sensitivity to remote homologs with fast runtime performance, and is able to detect homologs in the range of 10-30% sequence identity. Given the roughly 700 million years of evolutionary divergence between cnidarians and humans, we could not assume that homologs would be of higher sequence identity. In order to avoid confusion with other organisms, we built a custom HHblits database for *P. damicornis*, and one of its symbiotic algae, *Cladocopium goreaui (C1)*, formerly known as *Symbiodinium (S.) goreaui* (Clade C, type C1) (LaJeunesse et al., 2018) which we were able to query with human sequences of known function.

We know that corals adjust their behavior in response to external and internal cues, as illustrated by the following examples. Corals contract or extend their tentacles in response to light intensity (Levy et al., 2003). Coral larvae are known to prefer red over white surfaces for settling (Mason et al., 2011). Corals can fight, so they must be able to sense and attack the enemy or competitor (Yosef et al., 2020). Corals can distinguish organisms to tolerate (symbiosis) versus subjecting them to an immune attack to prevent disease (Mansfield & Gilmore, 2019). Corals prefer to eat plastic over copepods which may relate to a sense of taste (Allen et al., 2017). Since corals manage 90% of their energy from symbiotic algae (Allen et al., 2017; Wooldridge, 2010), they must also be able to measure and regulate nutrient balance. The key question that arises from these observations is: what are the molecular mechanisms underlying these behaviors?

Generally, signal sensing and response reactions in biological systems depend on membrane receptor signaling systems. Receptor activation involves the detection of the signaling molecule (ligand) outside the cell when the ligand binds to the receptor protein present on the surface of the cell (**Fig.1**). The signal transduction stage involves the activation of the receptor (conformational change) leading to the chain reaction of the activation of intracellular proteins. These signal transduction cascades trigger specific cellular responses. Thus, membrane receptors most generally are proteins that are coupled to intracellular signal transduction cascades. There are two types of membrane receptors, sometimes referred to as type I and type II receptors (**Fig.1**). Type I membrane receptors usually contain one or two transmembrane (TM) helices, and often carry enzymatic activity in their cytoplasmic domains, such as tyrosine kinase receptors like the epidermal growth factor receptor. Type II membrane receptors are G-protein coupled receptors (GPCR) which exclusively contain a seven-TM helical bundle. There are also ion channels which change their permeability in response to external signals, but are usually not classified as receptors (Nemecz et al., 2016). Membrane receptors have three major structural domains: extracellular (EC), transmembrane (TM), and cytoplasmic (CP). We will discuss one example in depth for each of the two types of membrane receptor proteins. For the type II membrane receptor family, we have chosen the GPCR sub-family of opsins. As an example for type I membrane receptors, we will discuss Toll-like receptors (TLRs).

**Figure 1.**
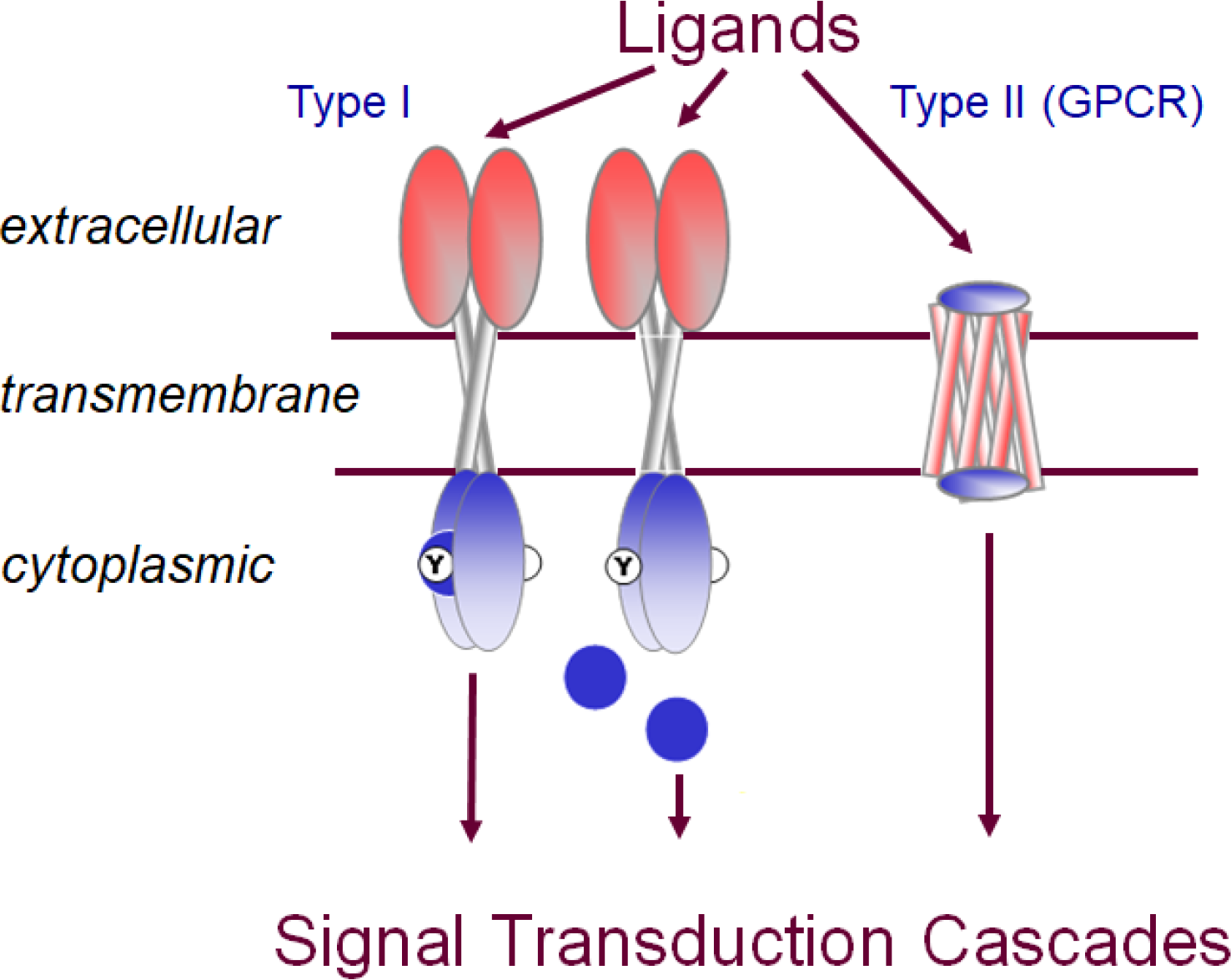
Schematic representation of type I and type II membrane receptors in signal transduction.

GPCRs respond to a large diversity of signals, from light in the opsin family to binding of hormones, neurotransmitters and even other proteins (Sakmar, 2010). Correspondingly, in humans, these receptors take a premier role as pharmaceutical targets and have thus been extensively studied (Sriram & Insel, 2018). Signal transduction by GPCRs is via binding and activation of heterotrimeric G proteins composed of alpha, beta and gamma subunits (Wettschureck & Offermanns, 2005). GPCRs are responsible for the majority of cellular responses to external stimuli. Upon activation by a ligand, the receptor promotes the exchange of GTP for GDP in its partner heterotrimeric G protein complex, leading to the dissociation of the alpha and beta gamma complexes from the receptor and each other. The G proteins then interact with other downstream effector proteins to mediate a cell response. In this study, we have used our remote homology detection pipeline to find human GPCR as well as G protein homologs in *P. damicornis* and *Cladocopium goreaui* (C1).

TLRs play a crucial role in innate immunity (Akira & Takeda, 2004). These membrane glycoprotein receptors recognize and respond to a variety of microbial (viral, fungal, and bacterial) components such as lipopeptides (Kang et al., 2009), peptidoglycans and lipopolysaccharides (Park et al., 2009), flagellin (Park et al., 2009; W. S. Song et al., 2017), DNA (Ohto et al., 2018), and RNA (W. Song et al., 2015). TLRs consist of leucine-rich repeats (LRR), and the Toll/interleukin-1 receptor (TIR) domain (Narayanan & Park, 2015). TLRs initiate signal transduction through interactions with TIR-domain containing adapters such as MyD88 (Dunne & O’Neill, 2005), that in turn recruits interleukin-1 receptor-associated kinase (IRAK) via interactions through death domains (Chen & Jiang, 2013). After phosphorylation, IRAK family proteins interact with the TRAF6 adapter. TRAF6 activates TAK1, a member of the mitogen-activated protein (MAP) kinase family (Dong et al., 2006), leading to the activation of NF-kB via kinase dependent signaling cascade involving IkB kinase complex (IKK-⍺,- , and -γ) and MAP kinases (ERK, p38, JNK), resulting in the expression of target genes (Tang et al., 2021). Viruses and bacteria are abundant in seawater and live in close association with corals (van Oppen & Blackall, 2019). The chemical crosstalk between corals and microbes plays an important role in coral growth and development. Corals need TLRs to communicate with microbes and it has been proposed that TLR signaling is conserved in corals (Williams et al., 2018). In the present study, we used our remote homology detection pipeline to find human TLR protein homologs in *P. damicornis* and *C. goreaui* (C1) to advance our understanding of coral innate immunity.

Using these two representative membrane receptor families, we will demonstrate that the stated problem of identifying remote homologues in non-model organisms is not just a remote sequence detection problem, but highlights the need for functional investigation of the putative proteins, which involves both systems and structural aspects of these proteins and their interaction partners. To this end, we focus on those positions in the alignments between model and non-model organism homologues that determine specificity of the respective protein’s functions. We will discuss below first the common elements of the pipeline, how we determine the pool of all membrane receptors in *P. damicornis,* how we subdivide this broad group into type I and type II membrane receptors, and how we refined these two multi-subfamily groups into functional classes by analysis of two of them, TLRs and opsins, by way of identifying the specificity determination positions, followed by analysis of the downstream signaling proteins.

## MATERIALS AND METHODS

### (a) Protein sequence retrieval

All non-*P. damicornis* and *C. goreaui* sequences were retrieved from the UniProt - Swiss-Prot Protein Knowledgebase, SIB Swiss Institute of Bioinformatics; Geneva, Switzerland (https://www.uniprot.org/). *P. damicornis* sequences (Cunning et al., 2018) were downloaded from http://pdam.reefgenomics.org/download/. *C. goreaui* sequences (Liu et al., 2018) were obtained from http://symbs.reefgenomics.org/download/.

Specific subgroups of sequences were identified and retrieved as follows and subjected to HHblits analysis as described in section (b).

#### (i) Human membrane receptor list

We have utilized two human membrane protein lists. The first one was published as the human membrane receptome in 2003 (Ben-Shlomo et al., 2003). We subjected the original list to an updated search in UniProt and retrieved 978 current UniProt entries, including GPCRs, provided in **supplementary file-S1.** The second was generated by using three alpha helix prediction tools followed by filtering of splice variants and clustering of the remaining genes (Almén et al., 2009). This list contains a total of 3,399 genes, including GPCRs, and is available from their supplement. The mapped *P. damicornis* entries (see below) are available in **supplementary file-S2.**

#### (ii) Human GPCR list

A comprehensive list of human GPCRs was extracted from an updated UniProt list of multi-species GPCRs (release: 2020_03 of 17-Jun-2020: 825 proteins (https://www.uniprot.org/docs/7tmrlist.txt). This list contained a total of 3093 GPCR sequences including 825 human GPCRs. See **supplementary file-S3** for the extracted human GPCRs. The mapped *P. damicornis* entries (see below) are provided in **supplementary file-S4.**

#### (iii) G protein list

Sequences of human G proteins were obtained from UniProt using keyword search, resulting in 16 alpha chains, 5 beta chains, and 12 gamma chains (Wettschureck & Offermanns, 2005). The complete list of these human proteins and their candidate *P. damicornis* homologues is provided in **supplementary file-S5.**

#### (iv) Toll-like receptor list

Protein sequences of TLRs of human (Akira & Takeda, 2004) and other organisms, namely Drosophila (Imler & Hoffmann, 2002), chicken (Nawab et al., 2019), frog (Ishii et al., 2007), and zebrafish (Kanwal et al., 2014), were retrieved from UniProt. TLR downstream signaling molecules (MyD88, TIRAM, TIRAP and TRIF) were also acquired from UniProt. The complete list with all the protein sequences used is provided as **supplementary file-S6**. The extracted *P. damicornis* TLR list is provided as **supplementary file-S7** (note there is a separate sheet for each organism).

### (b) Remote homology detection using HHblits

To enable organism specific searches, we created an online HHblits coral protein remote homology search tool (https://hhblits.cs.tufts.edu/), an installation of HHblits (Remmert et al., 2011) with genomic databases built for *P. damicornis* and *C. Goreaui*. Individual FASTA sequences were imported to this search tool and queries were run with an E-value cutoff of 10^-3^, single iteration, minimum probability of 20 (default), and with the minimum number of lines to show in the hit list expanded from 10 to 250. The jobID, email information, database information (e.g. *P. damicornis*) were submitted and the result output file was received by email or as batch predictions. Individual predictions contained the HHblits results of the submitted proteins in text format including protein sequence alignment, E-value, P-value, probability, column matched and score. In the case of GPCR, the result output was analyzed in a bidirectional fashion as follows. First, the top ranked *P. damicornis* hit was retrieved for each GPCR. Then a list of unique *P. damicornis* proteins were created from that, and the corresponding human GPCRs for which they appeared as top hits, provided as **supplementary file-S4.** For membrane receptors, the non-redundant lists of top ranked *P. damicornis* hits retrieved in the same way as for GPCRs are provided for both lists (**supplementary file-S2**). In the case of TLRs and G proteins, HHblits results were analyzed manually (see <u>Results</u>). For TLRs, top hits with 100% probability score were selected and analyzed separately for each model organism, i.e. human (Akira & Takeda, 2004), Drosophila (Imler & Hoffmann, 2002), chicken (Nawab et al., 2019), frog (Ishii et al., 2007), and zebrafish (Kanwal et al., 2014), before comparing them across organisms (Table 1). The list of *P. damicornis* hits for TLRs are provided as **supplementary file-S7.** A summary based on the frequency of occurrence of *P. damicornis* hits is provided in **Table 1**.

**Table 1.** Number of times a model organism shows homology to a given *P. damicornis* protein with 100% probability. Columns highlighted in green show at least one representation in each model organism studied.

| Organisms | *P. damicornis* proteins |  |  |  |  |  |  |  |  |  |  |  |  |
| --- | --- | --- | --- | --- | --- | --- | --- | --- | --- | --- | --- | --- | --- |
|  | 22934 | 22930 | 11599 | 9200 | 14109 | 17966 | 13021 | 15883 | 11734 | 21819 | 737 | 15877 | 9057 |
| Human | 5 | 5 | 10 | 7 | 4 | 3 | 3 | 2 | 2 | 2 | 1 | 1 | - |
| Zebrafish | 8 | 7 | 11 | 9 | 6 | 6 | 2 | - | - | 2 | 1 | - | - |
| Frog | 6 | 5 | 10 | 10 | 4 | 4 | 3 | - | - | - | - | - | 1 |
| Chicken | 7 | 7 | 9 | 9 | 2 | 2 | 1 | - | - | - | - | 1 | - |
| Drosophila | 9 | 9 | 7 | 9 | 2 | 3 | 2 | - | - | 1 | - | 1 | - |

### >(c) PROSITE analysis

The most highly ranked *P. damicornis* sequences retrieved through HHblits were subjected to PROSITE analysis (de Castro et al., 2006) to identify the presence of conserved domains (https://prosite.expasy.org/). Protein sequences were submitted to the PROSITE user interface and the results were analyzed and grouped by combining the domain schematic provided by PROSITE.

### (d) Transmembrane helix detection

In some cases, we verified the presence of a transmembrane region using TMHMM Server v. 2.0 (DTU bioinformatics, Department of Bio and Health Informatics) for the prediction of transmembrane helices (https://services.healthtech.dtu.dk/service.php?TMHMM-2.0).

### (d) Homology-modeling

Homology models were generated using Swiss Model (https://swissmodel.expasy.org/), an integrated web-based service dedicated to homology modelling of proteins (Waterhouse et al., 2018). We used the target-template alignment function of Swiss Model and provided the reconstruction of the full TLR5 (3J0A) combining crystallographic and cryo-electron microscopy data created by (Zhou et al., 2012) as a template to model *P. damicornis* 9200. We have also modelled our potential matches for opsins (629, 2270, 12246, and 19775) using squid rhodopsin as a template (2ZIY) (Shimamura et al., 2008). The models were evaluated for their global quality estimate and local quality score as per Swiss Model guidelines. The models were downloaded and analyzed using PyMOL (version-2.3.4, Schrodinger, LLC). These proteins were further structurally analyzed for ligand binding pocket and Ballesteros-Weinstein numbering system. Ligand binding pocket residues in GPCRs were extracted from previous chemogenomic analysis (Surgand et al., 2006).

### (e) Molecular docking studies of retinal with coral opsin homologs

Molecular docking was performed using retinal as the ligand with AutodockTools 1.5.6 (Harris et al., 2008). We standardized our docking experiments using squid rhodopsin with the following parameters: center_x=43.171, center_t=6.216, center_z=17.019, size_x=16, size_y=22, and size_x=26. Docking was performed by extraction after aligning the homology model with the squid rhodopsin space coordinates. The ‘exhaustiveness’ option was set as 32.0. The binding pocket was analysed using the Biovia discovery suite 2019 v19.1.0.18287 (Dassault Systemes Biovia Corp). Residues involved in the interactions for each model are listed in **supplementary Table-S1**.

### (f) Multiple Sequence Alignment of *P. damicornis*

Potential *P. damicornis* members of respective protein families were aligned to each other using MUSCLE (Edgar, 2004) (https://www.ebi.ac.uk/Tools/msa/muscle/) to examine if structurally relevant amino acids were conserved across family members. As an example, the alignment of opsin homologues in *P. damicornis* is shown in **supplementary Figure-S2**.

### (g) D-SCRIPT Analysis

We initially identified candidate homologs in *P. damicornis* for the human alpha, beta, and gamma G proteins using HHBlits (Remmert et al., 2011) (hhblits.cs.tufts.edu). We took the union of top hits to identify 124 candidate alpha proteins, 207 candidate beta proteins, and 5 candidate gamma proteins. We used the human pre-trained D-SCRIPT (Sledzieski et al., 2021) model to predict interaction between all pairs of alpha-beta, beta-gamma, and alpha-gamma subunits. We performed the same analysis in *Montipora capitata* (Shumaker et al., 2019), where we identified 184 candidate alpha proteins, 253 candidate beta proteins, and 4 candidate gamma proteins. We created a mapping between *P. damicornis* and *M. capitata* proteins using BLAST (Altschul et al., 1997) and identified best-bidirectional-hits, i.e. a pair *(P,M)* map to each other if *M* is the best BLAST hit in *M. capitata* for *P* and *vice versa*. We overlay the *P. damicornis* and *M. capitata* networks with each other using the mapping and identify as network-evidence candidate proteins where pairwise interaction is predicted between an alpha, beta, and gamma subunit forming a triangle in the network.

## RESULTS

### (a) Development of a pipeline: addressing the challenges with existing methods

An overview of the pipeline we developed to address the goal of identifying the repertoire of membrane receptors in the non-model organism, the coral *P. damicornis*, using known information from the model organism, human, is shown in **Figure 2**. The first step is to create a list of human proteins representing the function of interest, here membrane receptors. A list of membrane receptors in human had been published previously (Ben-Shlomo et al., 2003), but when retrieving the sequences from the UniProt database, several entries were no longer valid. We manually retrieved the updated UniProt ID’s by searching the database through protein names. The list of human membrane receptors obtained is available as **supplementary file-S1**. This list contains 978 human proteins. We also used another published list of membrane receptors (Almén et al., 2009), which contained 1352 human proteins reported using an outdated identifier format. Finally, a list of human GPCRs was extracted from a GPCR list in any organism from the UniProt database available at (https://www.uniprot.org/docs/7tmrlist.txt).

**Figure 2.**
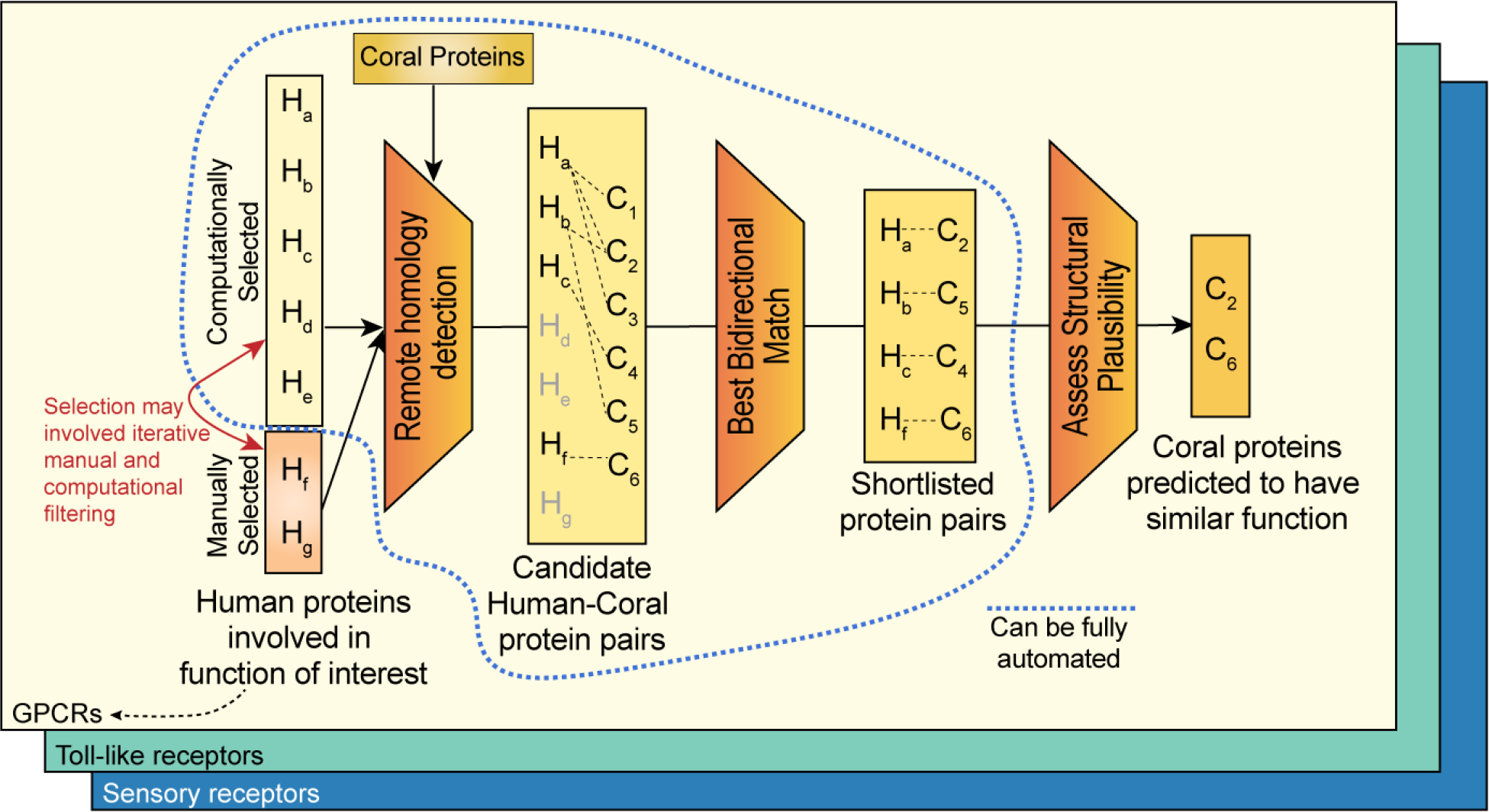
*In silico* protein prediction methodology.

Initially, we used BLAST, as well as several multiple sequence alignment (MSA) based tools to retrieve *P. damicornis* homologues, but after inspection of the results we concluded that the alignments between human and *P. damicornis* sequences were poor as judged by the number of gaps and the fractions of sequences aligned (data not shown). We concluded that due to the low sequence similarity between human and *P. damicornis*, we required a remote homology detection tool. HHblits is the accepted gold standard for remote homology detection (Remmert et al., 2011). Thus, all three human membrane receptor lists were searched against *P. damicornis* using our implementation of HHblits, available at https://hhblits.cs.tufts.edu/, where our implementation of HHblits contains genome-wide template libraries of HMM models of all the genes in several coral animal and symbiont genomes (principally *P. damicornis, M. capitata,* plus the clade C1 symbioint).

When mapping these three sets of lists to the *P. damicornis* sequences using HHblits, it became clear that the same genes mapped to multiple human query sequences. This is because there are many members of membrane receptor superfamilies, e.g. receptor tyrosine kinases or GPCRs in each organism itself already. We therefore created non-redundant lists of the top ranked *P. damicornis* sequences. We obtained 374 unique *P. damicornis* sequences from the 978 human plasma membrane receptome (older list), 329 from the 1352 human membrane proteome (newer list) and 151 from the 825 human GPCRs. These lists and their respective overlap is provided in **supplementary file-S2**. Combining the three lists and eliminating duplicates appearing in more than one of the lists yielded a total of 446 unique *P. damicornis* sequences. While all 151 GPCRs were found in both membrane receptor lists, there were 111 and 50 non-GPCR receptors missed if we considered only the older human receptor list as compared to the newer one. Thus, we conclude that there are 295 non-GPCR and 151 GPCR candidates in the *P. damicornis* membrane receptome.

It is important to note that, in the majority of cases, none of the quantitative parameters of the sequence alignment was able to differentiate between the hits. In other words, it was not possible to automatically assign a best human homologue for any given *P. damicornis* sequence. This is because different members of protein families have diverged in humans, and are more similar to each other, than they are to any *P. damicornis* sequence, which is evolutionarily most distant to all of them.

Given the fact that many more human sequences map to a smaller set of sequences in *P. damicornis* raises the question which functional sub-categories of these superfamilies are present in *P. damicornis*. Clearly, sequence alone is not sufficient to answer this question, and therefore identifying which of the functionalities of a given sub-family of membrane receptors is present in corals requires an analysis of known functional properties. This requires domain expertise and can no longer be fully automated in contrast to all previous steps (Figure 2). This final step of the pipeline is demonstrated for two examples below and necessarily branches according to the functions of the proteins of interest. The opsin subfamily was chosen as a representative example for the type II membrane receptor, the GPCR pipeline (section (b) below). The TLRs were chosen to represent the type I membrane receptors (section (c), below).

### (b) GPCR (type II receptor) pipeline branch

#### (i) Global analysis of GPCR families

The large GPCR family has been divided into subclasses based on a combination of pharmacological and sequence considerations, a classification which has been revised a number of times over the years (Surgand et al., 2006). Here, we use the clusters obtained with a chemogenomics approach based on the alignment of 30 critical GPCR positions supposed to face the ligand binding cavity (Surgand et al., 2006). Major classes besides olfactory receptors are Frizzled, Glutamate, Secretin and Adhesion families with the Class A rhodopsin family being split into 18 different clusters. When analyzing the human receptors for which *P. damicornis* sequences were found, these included chemokine, taste, glutamate, adrenergic, lipids, peptides, adenosine, amines, melanocortins, acids, chemoattractants, purines, frizzled, adhesion, prostanoids, and MAS-related receptors. Likely missing are melatonin receptors (with 2 low ranked options), vasopeptides, brain-gut peptides, SREB, secretin, opioid and glycoproteins. In all cases there are multiple human sequences that map to multiple (but less than humans) *P. damicornis* sequences. Detailed below is the evidence suggesting that *P. damicornis* can smell, taste and see light in sections (ii), (iii) and (iv), respectively.

#### (ii) Odorant receptors in *P. damicornis*

We found 105 human olfactory receptors that map to pdam_00017423-RA, 27 that map to pdam_00020860-RA, 17 that map to pdam_00017300-RA, while pdam_0005244-RA, pdam_0005376-RA and pdam_00016463-RA had one hit each. This suggests that there are at least 6 olfactory receptors in *P. damicornis*.

#### (iii) Taste receptors in *P. damicornis*

HHblits suggests that there are 15 remote homologs for the human taste receptors in the *P. damicornis* proteome: pdam_00004028-RA, pdam_00009436-RA, pdam_00000629-RA, pdam_00002659-RA, pdam_00013619-RA, pdam_00021435-RA, pdam_00022798-RA, pdam_00017219-RA, pdam_00001145-RA, pdam_00013621-RA, pdam_00010275-RA, pdam_00004281-RA, pdam_00002512-RA, pdam_00020500-RA, and pdam_00016973-RA. Transmembrane helix prediction using TMHMM confirmed the presence of 7 transmembrane helices in 13 out of 15 proteins. pdam_00002512-RA, and pdam_00009436-RA have more than 7 predicted transmembrane helices.

#### (iv) Vision receptors (opsins) in *P. damicornis*

The HHblits result showed that the 11 members of the human opsin family mapped to 7 *P. damicornis* sequences. However, there were other members of the Class A GPCR family that also mapped to the same *P. damicornis* sequences. To find out which of the HHblits remote homology predictions were likely true representations of visual functions, we analyzed the ligand binding pockets of these proteins. Light detection is the main function of rhodopsin. Therefore, the homologous proteins must have an ability to bind with a light sensitive retinal ligand and should possess a conserved ligand binding pocket. This pocket should show sequence similarity with human opsin protein. Considering this hypothesis, we analyzed the human opsin ligand binding pocket and compared our analysis with *P. damicornis* opsin homologues. We obtained the profiles of 30 cavity-facing amino acids used for clustering of the human GPCRs including opsin (Surgand et al., 2006). Thus, the approach outlined here for opsins can be extended to other GPCRs. The first step was to verify the presence or absence of these amino acids important for ligand binding in the *P. damicornis* sequences for the opsin family. The 30 residues are spread out throughout the sequence, and the Ballesteros-Weinstein numbering scheme used to identify aligned positions in GPCRs clearly showed that these residues are located across the 7 transmembrane helices. We compared these 30 residues against the top hit for OPSD and OPN5, and the result showed that only 11 out of 30 residues are the same between OPSD and their *P. damicornis* homologue.

Further, we had previously determined the minimal ligand binding pocket that is capable of exerting the function of a GPCR: transmission of the ligand binding signal to the downstream signaling proteins (Moitra et al., 2012). The approach used GREMLIN, Generative Regularized Models of Proteins, to identify long-range interactions from co-varying amino acid positions in multiple sequence alignments (Balakrishnan et al., 2011). GREMLIN learns an undirected probabilistic graphical model known as a Markov Random Field (MRF). Unlike HMMs, which are also graphical models, MRFs can model long-range couplings (non-sequential residues). We performed GREMLIN analysis on GPCRs, statistically evaluated different sizes of ligand binding pockets and found that a pocket as small as 4 residues still shows significant enrichment of edges over null. This means that four residues connect maximally to the rest of the protein orchestrating the global conformational change inherent to GPCR function and are required for signal transduction. Our analysis showed that 3 of these 4 residues are conserved across human opsin and the *P. damicornis* homologue. This is strong evidence that this protein is a functional GPCR.

Finally, GPCRs involved in vision require a lysine in a specific location of the ligand binding pocket to covalently attach via a Schiff base. The Schiff base is stabilized by another residue in the sequence, that is glutamic acid (E113), forming a counter ion to lysine K296, (highlighted by red arrows in **Figure 5**). We can see that the *P. damicornis* homologue contains the lysine required for retinal binding, but has a tyrosine instead of glutamic acid in the counter ion position. This substitution is expected to cause a shift in the absorbance maximum but not a loss of function. Thus, we can conclude that this *P. damicornis* sequence indeed represents a functional opsin protein, likely with an absorbance maximum different from that of rhodopsin and more like the OPN5 protein, which also has a tyrosine at this crucial position. Repeating this process for all of the *P. damicornis* homologues allowed us to eliminate proteins that did not fit the requirements of GPCR action or retinal binding function and narrow down the most closely related opsin protein. We predict that there are 4 *P. damicornis* opsin proteins (in contrast to 11 human opsin proteins) and these are most similar to OPN5, OPSG, OPSX and OPSR. We conclude that *P. damicornis* can likely sense and respond to light and even distinguish colors. Previous research showed that the larvae preferentially settle on red as opposed to white surfaces (Foster & Gilmour, 2016). Consistent with this behavioral finding, is the inclusion of the red opsin protein (OPSR) in our list.

#### (v) Homology modeling and molecular docking studies with opsin homologs

To confirm the structure and functional relationship of rhodopsin with putative coral rhodoposin homologs, we have performed structural modeling of the top coral rhodopsin hits using the squid rhodopsin structure as a template (**Figure 6**). We carried out molecular docking to these homology models that suggested that the retinal was indeed able to bind with the putative ligand binding pocket in the coral rhodopsin homologous structures. This suggests that these are strong candidates to be considered as coral rhodopsin homologs and may play an important role in light sensing mechanisms. We have summarized the details of the retinal interactions with squid rhodopsin and coral rhodopsin homologs in **Supplementary Table 1.**

### (c) GPCR binding partner: G Proteins

To gather further evidence that the proteins identified were functional opsins, we investigated their function, namely binding and activation of the G proteins upon ligand binding (or light activation of the retinal ligand in the case of opsins). G proteins are heterotrimeric proteins consisting of alpha-, beta- and gamma-domains. There are 16 different alpha chains, 5 beta chains, and 12 gamma chains in humans (Wettschureck & Offermanns, 2005), and their UniProt ID’s are provided in **Supplementary File S5, tab 1**. HHblits search of the alpha and gamma chains yielded six potential *P. damicornis* alpha chains and five potential gamma chains, see **Supplementary File S5, tab 2**. We were unable to retrieve potential beta chains reliably, due to the beta propeller structure of the beta chains being a very common structural motif found across multiple protein families.

#### (i) Multiple Sequence Alignment of *P. damicornis* alpha chains

The potential *P. damicornis* alpha chains were aligned with the human G-alpha subunit that binds to rhodopsin, called transducin, with gene symbol GNAT1 using MUSCLE (see <u>Methods</u>) to examine if structurally relevant amino acids were conserved. Amino acids of GNAT1 which interact with rhodopsin, GNB1, and GTP were identified from co-crystal structures (PDB ID: 6OY9, 1TAD) by selecting residues within 5 Å of the relevant structure in PyMOL. These residues are highlighted in **Figure 6A** using boxes colored orange, blue, and green, respectively. The alignment shows that the GTP binding pocket residues are the most conserved, the receptor binding residues are the least conserved, and the beta chain binding residues are more conserved than the receptor binding residues but less conserved than the GTP binding residues. This is to be expected, as the GTP pocket will need to bind to the same structure in corals as it does in humans, whereas the receptor and beta chain binding residues will need to bind to coral receptor and beta chain homologues, which will be structurally different. Because the conservation patterns are similar across all 5 *P. damicornis* G-alpha candidates, we are not able to select a top candidate from this pool that is most likely representative of the rhodopsin-binding G-alpha subunit, transducin (GNAT1).

#### (ii) Network analysis of putative *P. damicornis* G protein subunits

To overcome the challenge that sequence alignment alone was not sufficient to narrow down the choices of *P. damicornis* sequences related to G proteins, we leveraged computational prediction of protein-protein interactions (PPI) to characterize the relative likelihood of the top *P. damicornis* hits being truly involved in GPCR binding activity. Applying D-SCRIPT (Sledzieski et al., 2021), a recently introduced sequence-based deep learning model for PPI prediction, we performed two complementary PPI analyses. In the first, we performed an all-vs-all computational screen of interaction between all the candidate G-alpha, beta and gamma proteins in *P. damicornis*, reasoning that the true positive hits will display the expected interaction patterns of alpha-beta and beta-gamma binding. Our predicted PPIs broadly conformed to this expectation, with pdam_00017900-RA, pdam_00011071-RA, pdam_00023984-RA, pdam_00007710-RA, pdam_00014456-RA, pdam_00011840-RA being the strongest candidates for alpha subunits, pdam_00000168-RA the strongest candidate for a beta subunit, and pdam_00000526-RA the strongest candidate for a gamma protein. To gain further confidence in our estimates, we also performed a similar screen of G protein subunits in *M. capitata* (**Figure 6C**) and assessed if homologous pairs across the two coral species (estimated by a bidirectional best hit analysis) had similar PPI patterns (see <u>Methods</u>). These results further supported our estimate, as the same connectivity patterns observed in *P. damicornis* are also observed between homologues in *M. capitata*.

#### (iii) Conservation of structural interfaces in G protein complexes

To further confirm the predicted G protein complex compositions, D-SCRIPT was then used to assess the structural plausibility of individual G-alpha candidates in *P. damicornis.* We performed an *in silico* mutagenesis study of each G-alpha candidate, evaluating these mutations by how they changed the D-SCRIPT score for interaction with candidate G-beta proteins pdam_00000168-RA and pdam_00014586-RA (**Figure 7**). Reasoning that a conservative test would be to require the same binding mechanism as seen in human G-alpha-beta interaction, we first aligned candidate G-alpha protein sequences in *P. damicornis* against human G-alpha proteins GNAT1, and used the multiple sequence alignment to identify the residues in coral G-alpha where the corresponding location in GNAT1 is known to play a role in binding to G-beta proteins. We tested if random *in silico* mutations at these locations had a particularly deleterious effect on the likelihood of PPIs with coral G-beta candidates (average of 50 trials). We compared these likelihoods to the original predicted probability of interaction, and the predicted probability of interaction after *in silico* mutations at random sites (average of 50 trials). Two candidates, pdam_00014456-RA and pdam_00011071-RA, showed sharply decreased likelihood of a PPI occurring specifically when the sequences were mutated at putative binding locations -- suggesting a binding mechanism similar to human G-alpha-beta interaction is conserved in determining the interaction likelihood of coral G-alpha-betas.

### (d) Toll-like receptor (TLR, type I receptor) pipeline branch

#### (i) TLRs in *P. damicornis*

After having demonstrated that there are functional GPCR (aka type II receptor) proteins and pathways in *P. damicornis*, we next investigated a representative of the type I receptors, namely TLRs. Because there are large differences in the numbers of TLRs in different organisms, we retrieved TLR sequences not only from humans (Akira & Takeda, 2004), but also zebrafish (Akira & Takeda, 2004; Kanwal et al., 2014), frog (Ishii et al., 2007), chicken (Nawab et al., 2019) and Drosophila (Toll proteins) (Valanne et al., 2011). After subjecting these sequences to HHblits prediction, we extracted all coral proteins that showed a minimum of 100% probability, and grouped them by their homology to a given organism. The results are summarized in **Table 1**. *P. damicornis* proteins 22934, 22930, 11599, 9200, 14109, 17966, and 13021 were observed as homologues to at least one TLR from each of the five model organisms studied. Because many type I receptors are often composed of multiple domains, we then subjected these proteins to PROSITE analysis in addition to homology modeling.

#### (ii) PROSITE analysis showed similar domain signatures in *P. damicornis* proteins

PROSITE analysis was used to identify domain structures within putative TLRs. **Figure 3A** illustrates that in human TLRs, there are multiple copies of leucine rich repeat (LRR) domains present on the extracellular side of the receptor and a Toll/interleukin-1 (IL-1) receptor (TIR) domain in the intracellular side in human TLRs. In contrast, *P. damicornis* 22934, 22930, 15883, 15877, 11734, and 11599 were devoid of LRRs and only displayed the TIR domain. This suggests that these proteins may not belong to the typical TLR family. On the other hand, PROSITE analysis of *P. damicornis* 14109 and 17966 homologues showed a large number of LRRs, plus a cadherin domain, EGF_CA (Calcium-binding EGF domain), Thrombospondin, type 3 repeat (TSP3), Thrombospondin C-terminal domain profile (TSP_CTER) domains on the extracellular side and the TIR domain on the intracellular side. Because of these additional domains, it is possible that these are TLRs but with different or additional functions as compared to their endosomal human counterparts. A third domain composition was revealed by PROSITE analysis of *P. damicornis* 737 which only had LRRs and no TIR domain. Therefore, this homologue was rejected as a TLR candidate. The only domain composition similar to that of human TLRs was observed for *P. damicornis* homologue 9200, which included multiple LRRs in the extracellular domain and a TIR domain in the cytoplasmic domain. For further analysis, we selected 9200 for homology modeling due to its matching profile with human TLRs.

**Figure 3.**
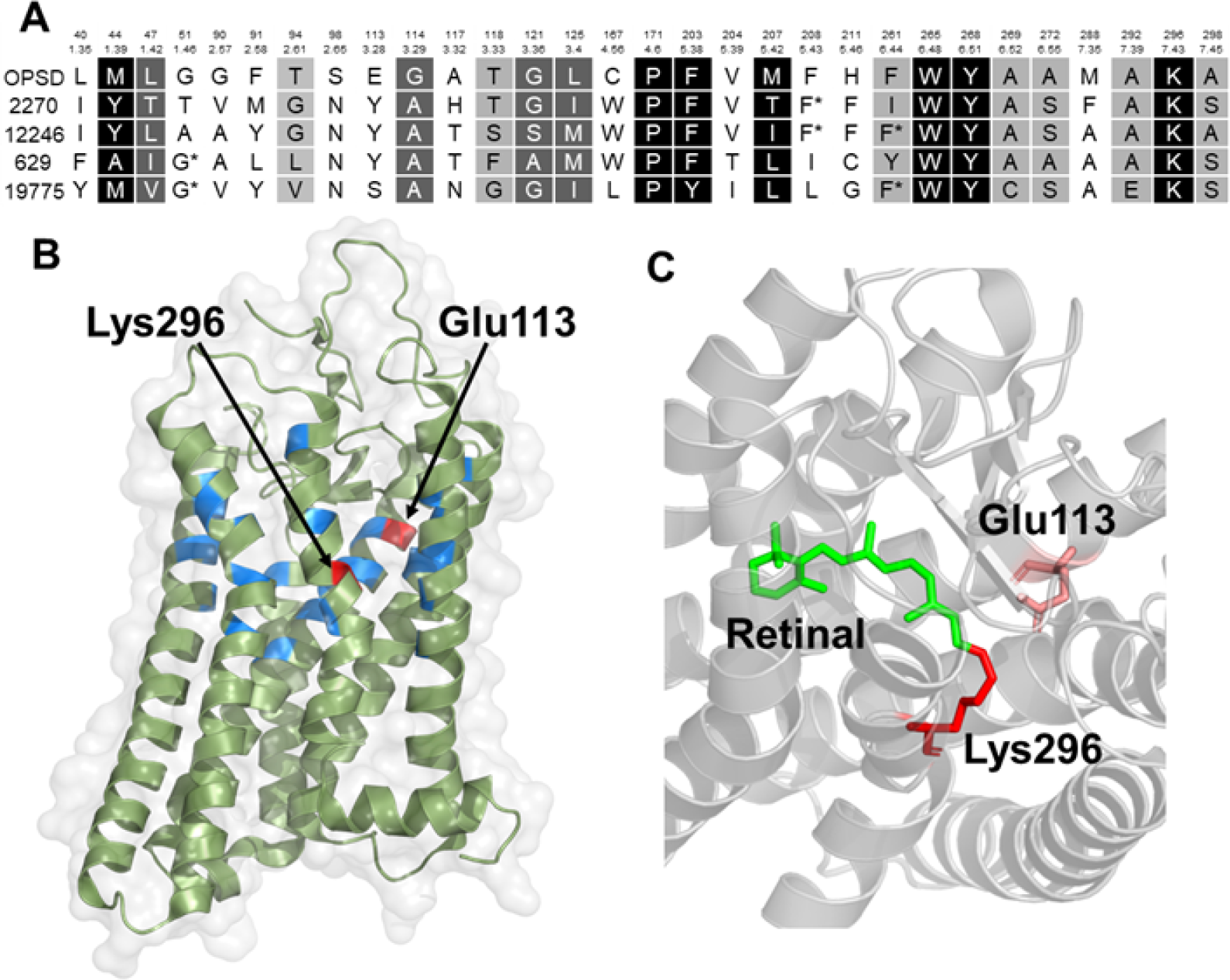
Sequence alignment and active site analysis of rhodopsin. (A) Alignment of rhodopsin and coral proteins and scoring based on Ballesteros-Weinstein (colored according to the degree of similarity: white foreground/black background, 100%; white foreground/grey background, > 80%, black foreground/grey background, > 60%). (B) Active site of bovine rhodopsin showing the residues involved in active site. (C) Active site showing the interaction of retinal with Lys296 and Glu113.

#### (iii) Structural similarities between human TLR5 and *P. damicornis* TLR

Swiss Model matching for query *P. damicornis* protein 9200 retrieved human TLR5 (3j0a) as the top hit. We refined the model by removing several missing regions from the coral TLR model (**Figure 4**). Using TMHMM with 9200, we identified a transmembrane helix region (649 -671) between the extracellular and the intracellular region, supporting the organization as a typical type I receptor.

**Figure 4.**
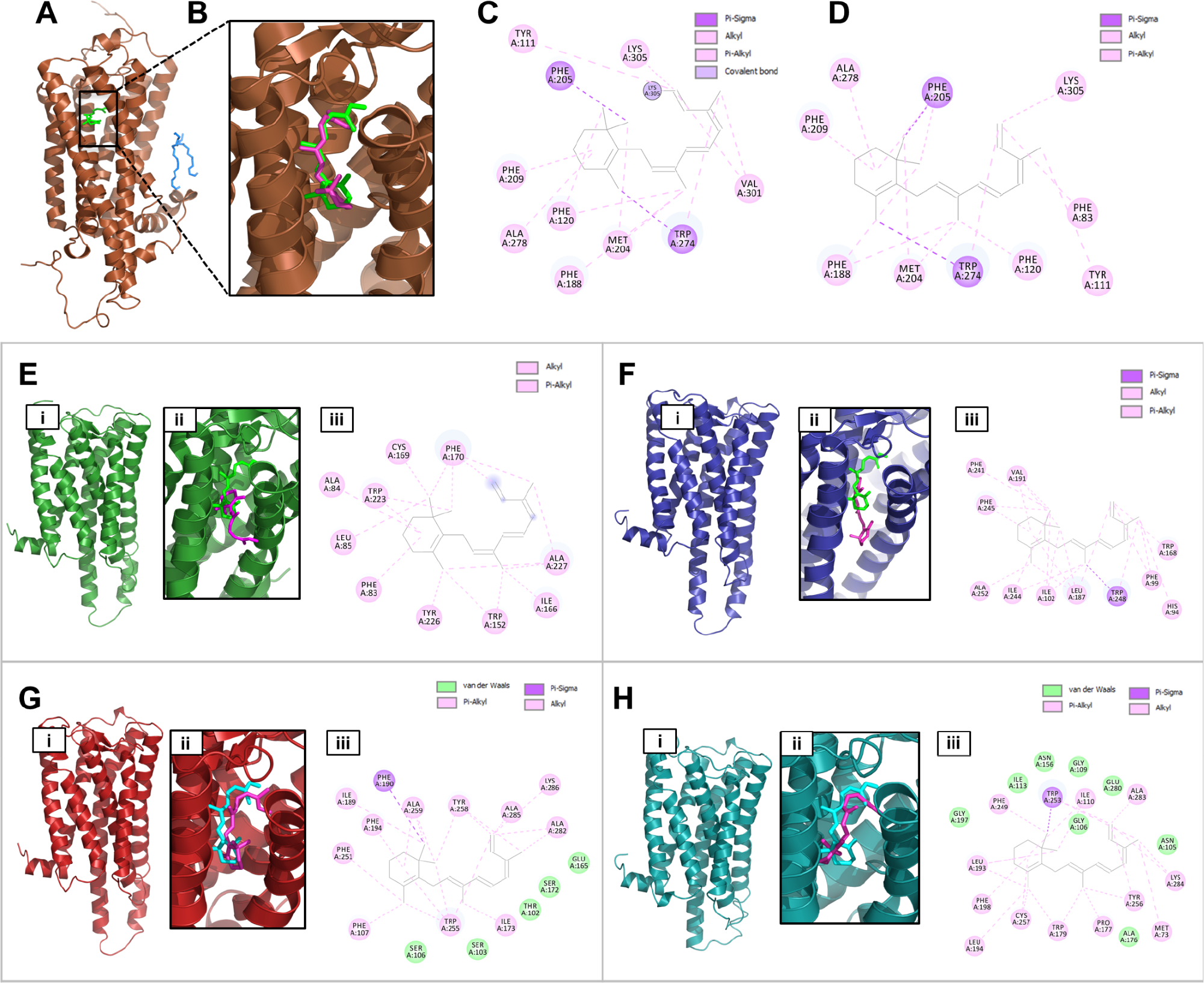
Homology modeling and molecular docking of rhodopsin receptors. (A) Crystal structure of squid rhodopsin (2ZIY). (B) Retinal in crystal structure and the docked conformation. (C, D) Molecular interactions of retinal in crystal structure vs. docked confirmation. (E, F, G, H) Homology modeling and docking studies with 629, 2270, and 19775, (i) homology model, (ii) retinal confirmations: natural (green) vs. docking pose (magenta), (iii) molecular interaction of docking pose (details of interactions and the docking scores are provided in **supplementary table 2**).

**Figure 5.**
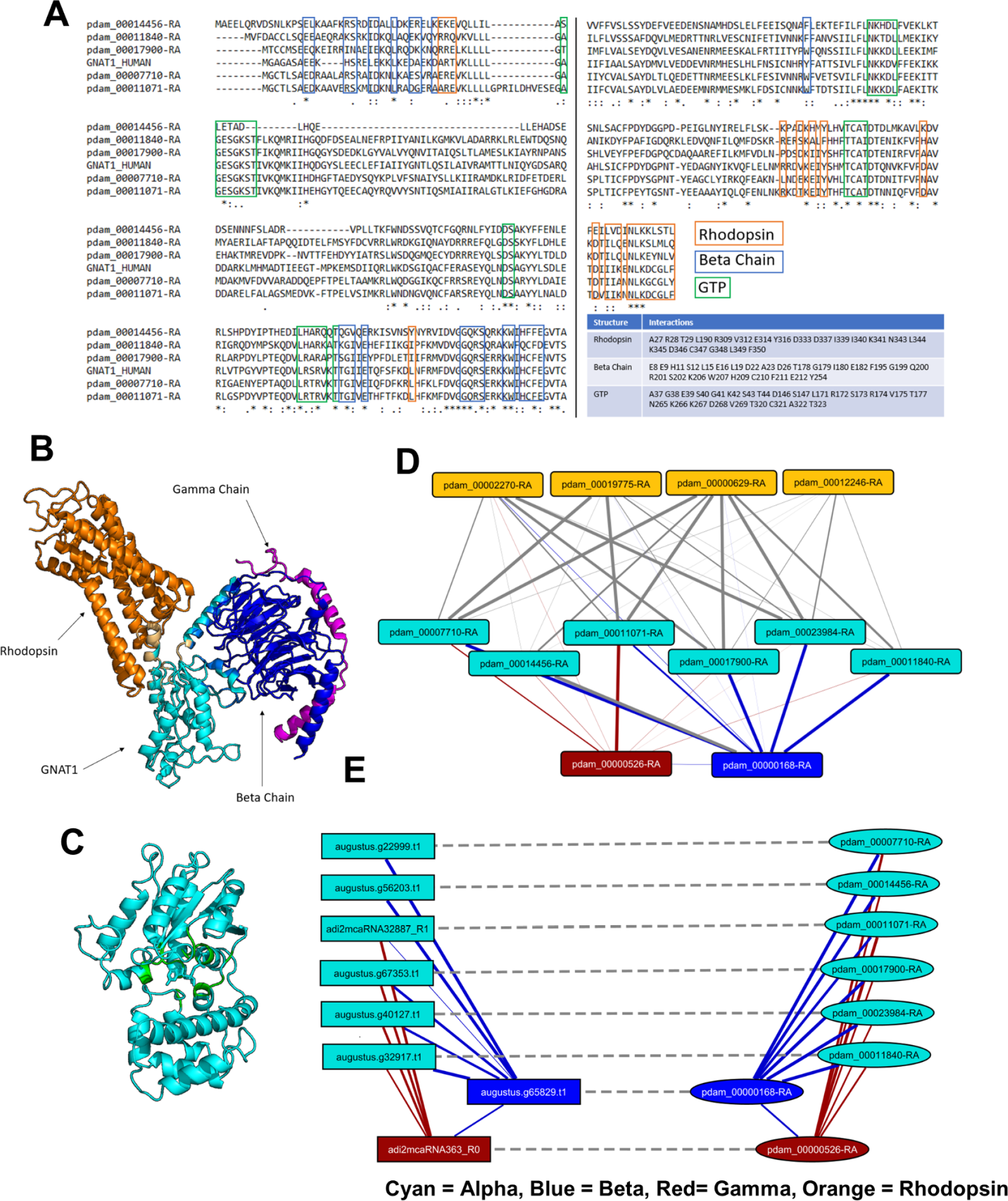
(A) MSA of GNAT1 with coral homologues and crystal structures of GNAT1. Amino acids of GNAT1 which interact with rhodopsin, GNB1 (beta chain), and GTP are boxed in blue, red, and green, respectively. The same amino acids are also listed in the table. In the crystal structures, GNAT1 (cyan) binding interfaces with rhodopsin (orange) and the beta chain (blue) are shown in (B) and the GTP binding pocket (green) is shown in (C). (D) Predicted interactions in *P. damicornis* between alpha (cyan), beta (blue), gamma (orange) chains and rhodopsin (orange). (E) Predicted candidate G protein interactions in *P. damicornis*, and the predicted interactions of the corresponding candidates in *M. capitata* (solid line: predicted interaction; dashed line: best-bidirectional BLAST hit).

**Figure 6.**
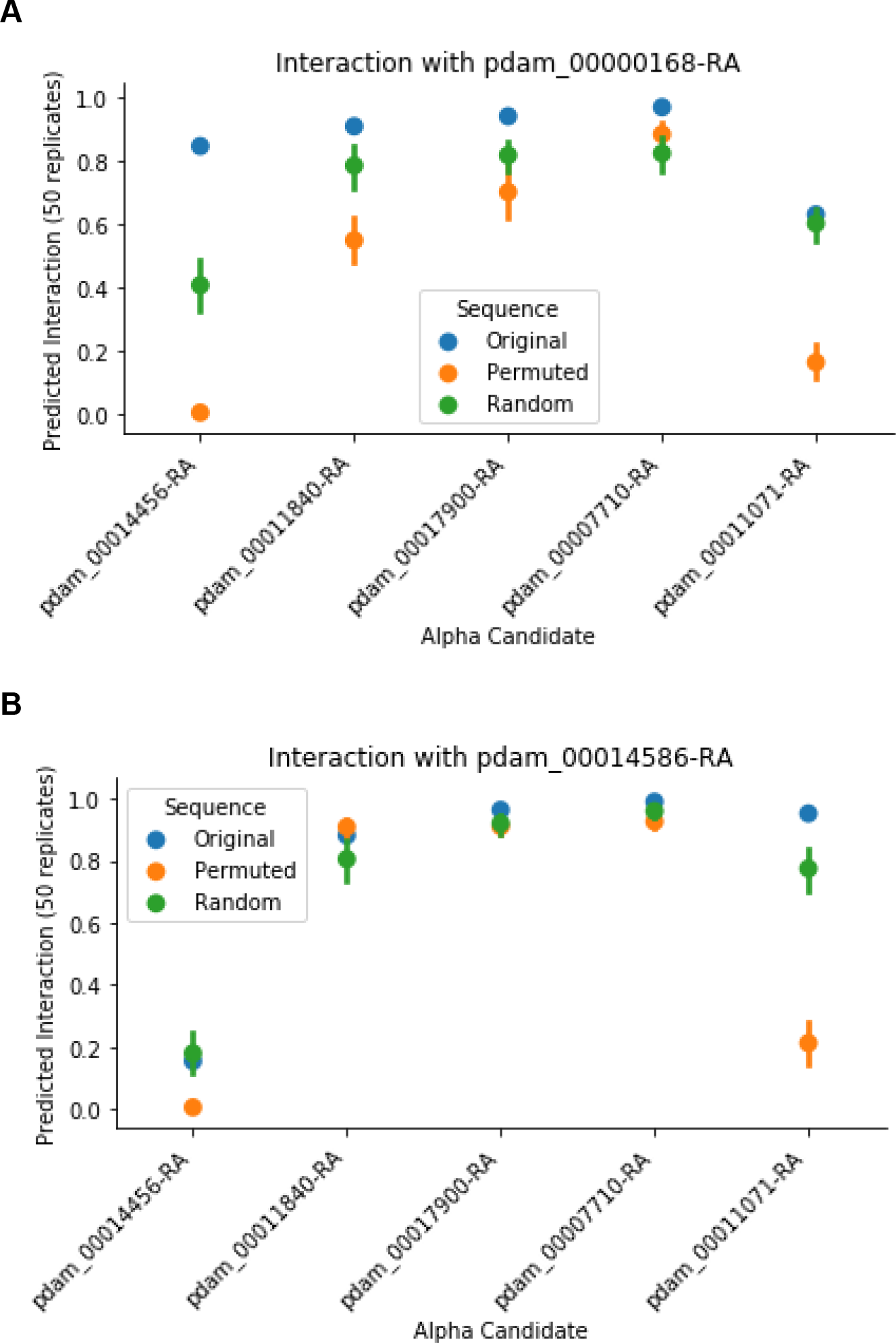
Predicted interaction of candidate alpha proteins with beta candidate proteins `pdam_00000168-RA` (A), `pdam_00014586-RA` (B). *Original:* Original predicted probability. *Permuted:* Average predicted probability of 50 samples with 25 beta-binding residues randomly perturbed. *Random*: Average predicted probability of 50 samples with 25 randomly chosen residues randomly perturbed.

#### (iv) MyD88 homologue as the possible downstream partner

TLRs utilize multiple adapter proteins to transmit the signal to the inside of the cell, namely MyD88, TIRAM, TIRAP and TRIF adaptor proteins. Our HHblits results of these proteins showed presence of a coral homologue only for MyD88.

## Discussion

We present a general pipeline applicable to any non-model organism to explore functions based on detectable homology to critical proteins in humans or other model organisms. The strategy involves the following steps (Figure 2). The first step is to create a list of related proteins representing a function of interest such as membrane receptors, or ion channels, or proteins involved in wound healing etc. Often, we can use computational expansion of proteins we know represent this function, but depending on the complexity and relatedness of the group of proteins, it may require manual selection. Once a suitable group of proteins has been established, we subject this list of sequences to remote homology detection, using HHblits. This usually yields a large collection of putative distant homologs. Traditionally, you would choose the top ranked hit and assume that this hit represents the best match to the query’s function. In the case of evolutionarily distant species, this approach fails because a protein family may have already diversified within each individual organism. There can be a smaller or a larger number of family members in one organism as compared to the other organism, and this may represent presence/absence of a particular function or sub-function. This creates the challenge that we need to distinguish which remote homologues in fact will have conserved the function of interest of the original query. The next step is therefore to identify the best bidirectional match from the candidate protein pairs to create a shortlist. So far, these steps can all be fully automated. Next, however, we need to assess the structural plausibility of each possible match, and this requires manual investigation that depends on the availability of structures and expert knowledge on the function of interest. We can think of this as casting a very wide initial net based on recovering a weak signal of sequence similarity, and then using 3D structural models to focus attention on a particular small number of functionally important residues that can provide a signature for function conservation. This sequence signature is based on a protein-protein interaction or a protein-ligand interface. We have presented several tailor-made approaches to this challenge. In the case of GPCRs, we looked at specificity determining positions (Capra & Singh, 2007; Kalinina et al., 2004), which can then be confirmed by doing 3D structural modeling, to check that the active sites and binding pockets that are expected, should the functional role of the protein be conserved, are indeed present. In the case of G proteins, we derived function based on protein-protein interactions, where we have used our recent deep learning method (Sledzieski et al., 2021) to perform *in silico* mutagenesis studies to further help us distinguish sequences which are likely to allow us to correctly transfer functional annotation from their human homologs from those that may have other functions. In the case of TLRs, we have used PROSITE to identify presence or absence of entire domains within the sequences, and used expert knowledge to evaluate if the location of these domains matches the functional expectation (e.g. if the domain faces the inside or outside of the cell). To this end, we compared PROSITE domain family predictions for the human query proteins and the putative coral homologues. All human TLR showed the presence of LRRs in the extracellular domain, and TIR domain in the cytoplasmic region. In contrast, in the 13 putative coral homologues, only one coral sequence of these candidates shows the expected (see Figure 1, type I receptor organization) extracellular LRR and intracellular TIR domain, separated by a transmembrane helix. This finding allows us to narrow down the candidate list to a single sequence.

The availability of three-dimensional structures for one or more members of the protein family of interest opens the possibility for in-depth structure modeling using homology. Although the quality of homology models depends on the degree of sequence conservation, structurally aligning remote sequences goes a long way in interpretation of their structure conservation as long as we can have confidence in the alignment. For example, despite near negligible sequence conservation between rhodopsin and metabotropic glutamate receptors, the most distant members of the GPCR family, we were able to predict the pharmacological outcome of ligand binding based on structural alignment (Yanamala et al., 2008). With the continuous improvements in *de novo* structure prediction (Jumper et al., 2021), this task will become more and more feasible even for proteins that do not have any homologous structures available, i.e. for which no structural template can be found.

One challenge not currently addressed by our pipeline is the identification of *de novo* proteins with *de novo* functions. While homology modeling allows us to detect proteins that are present (with mutations) in both species, we acknowledge that just as some proteins novelly evolved in vertebrates and are not present in cnidarians, there is the possibility of complexes that novelly evolved in corals, which this approach cannot detect.

The results of our study have several biological implications. Cnidarians as evolutionarily early animals offer great opportunities to study the evolution of important biological functions, such as sensing and signaling, and understanding host-microbe communication. TLRs are used for molecular communication with microbes. These receptors interact with specific microbial antigens and play a major role in innate immunity. We obtained strong evidence for TLR presence in *P. damicornis,* in line with several previous reports suggesting the presence of TLR mediated signaling in corals. Analysis of a cnidarian genome indicated the presence of immune related genes and the presence of Toll/TLR pathways, membrane attack proteins, and complement pathway associated signaling molecules (Miller et al., 2007; Nyholm & Graf, 2012). RNA-Seq data also supports the ability for immune responses in corals (Anderson et al., 2016; Cunning et al., 2018). One study showed that the muramyl dipeptide (MDP), a bacterial cell wall component, was found to trigger the up-regulation of GTPases of immunity-associated proteins in *Acropora millepora* (Weiss et al., 2013). Finally, the transcriptomic expression during thermal stress and pathogen challenge with *Vibrio coralliilyticus* in *P. damicornis* also supports the regulation of Toll/TLR and prophenoloxidase and complement pathways (Vidal-Dupiol et al., 2011). We now add detailed protein structural analysis to these reports, and also provide evidence that *P. damicornis* carries the gene for the TLR adapter protein, MyD88 (*P. damicornis* id 15711).

Of particular note is that our results suggest a much smaller number of proteins involved in TLR signaling than in the model organisms human, chicken, zebrafish, frog, and Drosophila. We predict that there may only be a single, unique TLR homolog in the coral *P. damicornis* that exhibits all the features expected for TLRs, and is most similar to TLR5 in human **(Figures 7,8).** For comparison, **Table 1** shows the frequency of each *P. damicornis* protein for a specific TLR protein in various model organisms. Furthermore, we have found only one (MyD88) of the four known adapter proteins MyD88, TIRAM, TIRAP and TRIF. This suggests the presence of a less diversified and simpler TLR signaling pathway in coral as compared to higher eukaryotes.

**Figure 7.**
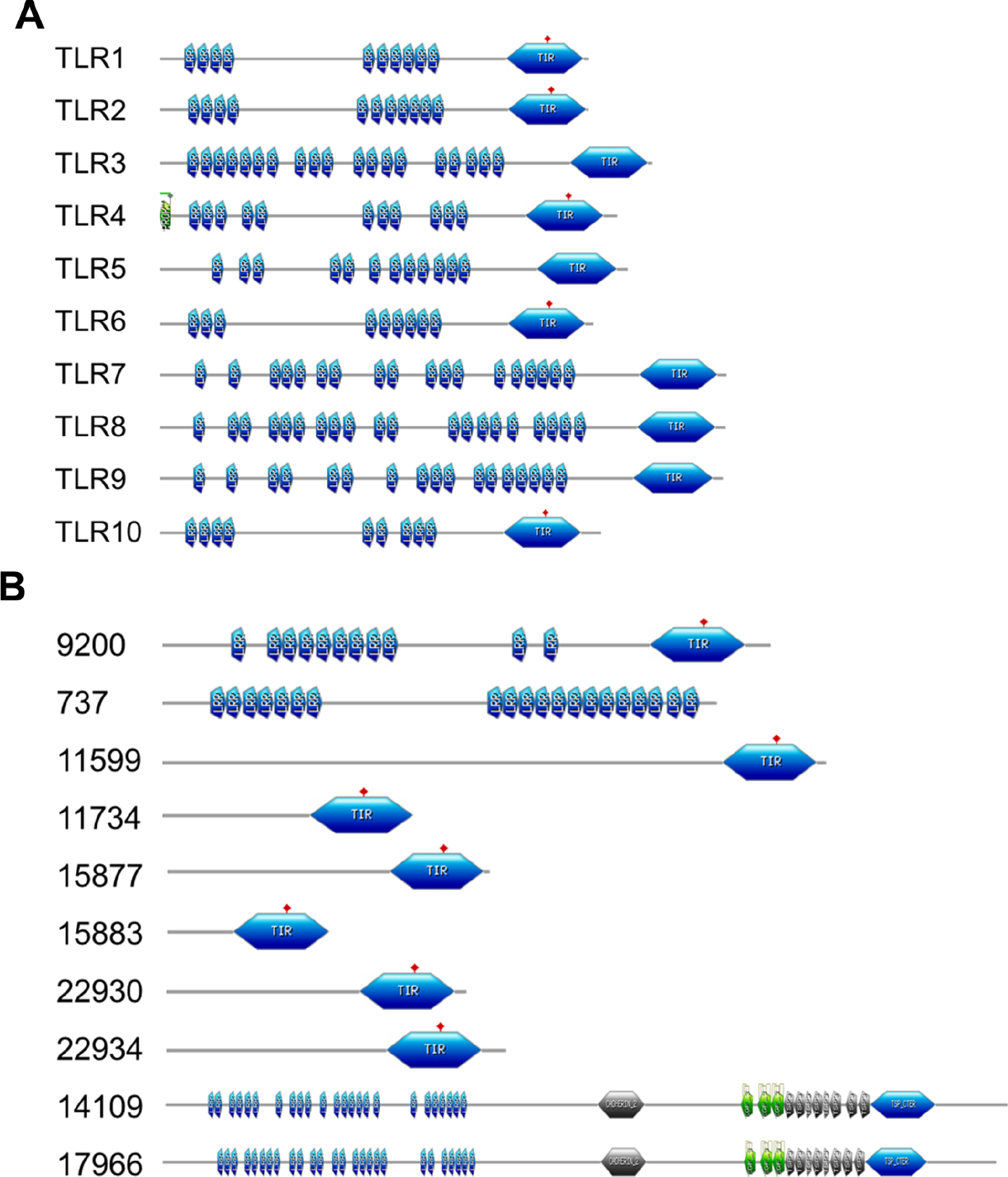
PROSITE analysis of human and coral TLRs. (A) PROSITE domain mapping of human TLRs (B) domain mapping of possible *P. damicornis* TLRs homologues.

**Figure 8.**
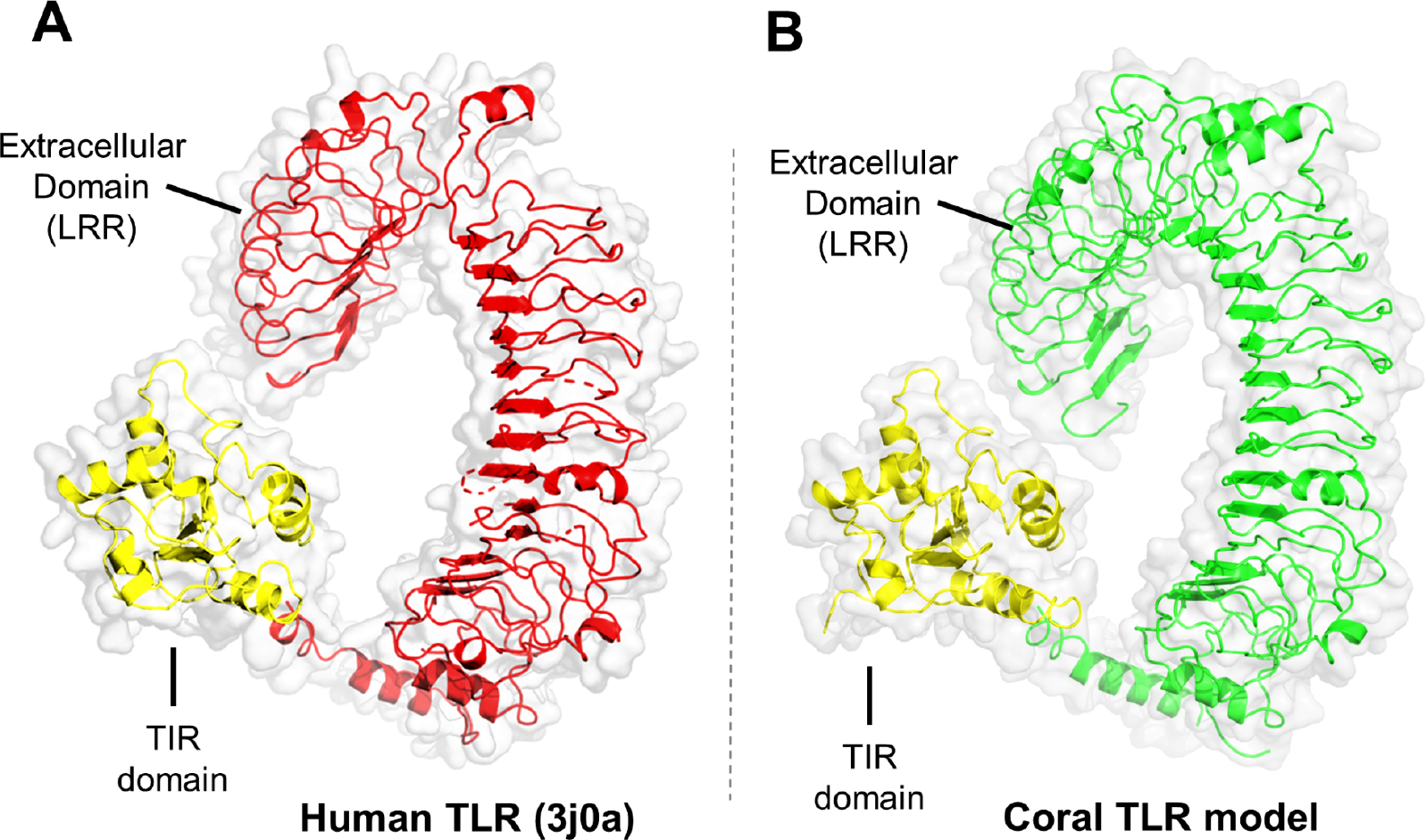
Coral TLR model construction using full-length TLR5 homology model structure. (LRR: leucine rich repeats). (A) homology model of full-length human TLR5 protein (3j0a) (B) model of *P. damicornis* 9200 protein using human TLR5 homology model as a template.

A similar conclusion was reached in the analysis of the large and diverse GPCR family. The human genome alone encodes 825 GPCRs, while we only found 151 GPCRs in *P. damicornis*. This is also reflected in the analysis of the opsin subfamily which has 11 members in human and we predict 4 in *P. damicornis*. Looking at reports of GPCR family size in evolutionarily early organisms suggest that there is a dramatic expansion in the numbers of GPCR when transitioning from unicellular to multicellular organisms. For example, choanoflaggelata have only 10 GPCR (Hake, 2019), while the Placozoan Trichoplax adhaerens has 420 (Srivastava et al., 2008), while sponges have 330 (Krishnan et al., 2014), Nematostella vectensis corals have 890 (Schiöth et al., 2010) and hydra has 1200 (!) (Chapman et al., 2010). Evolutionary analysis of the sub-families suggests that the doubling that takes place between Trchoplax adhaerens and Nematostella vectensis is roughly maintained and most sub-families remain relatively constant in their size distributions, with the Class A rhodopsin-like family being the largest in all species (Jékely, 2013; Nordström et al., 2011).

Taken together, both TLR and GPCR analysis suggest that *P. damicornis* represents a transition species that carries the minimal number of essential components for signaling and innate immunity, while additional functionality and fine tuning is achieved through diversification of this small pool of proteins in higher evolved organisms. Thus, being able to differentiate between early basic versions of a given function and finding the reasons for the need to diverge emphasizes the utility of this method assuming the best hit is the most functionally relevant hit. This may help to identify possible functions for currently not conclusively annotated proteins in *P. damicornis* and other non-model organisms.

## Acknowledgments

This work was sponsored by NSF grants HDR: DIRSE-IL 1940169 and RAPID 2031614.

## Supplementary Figures and Tables

**Figure S1.**
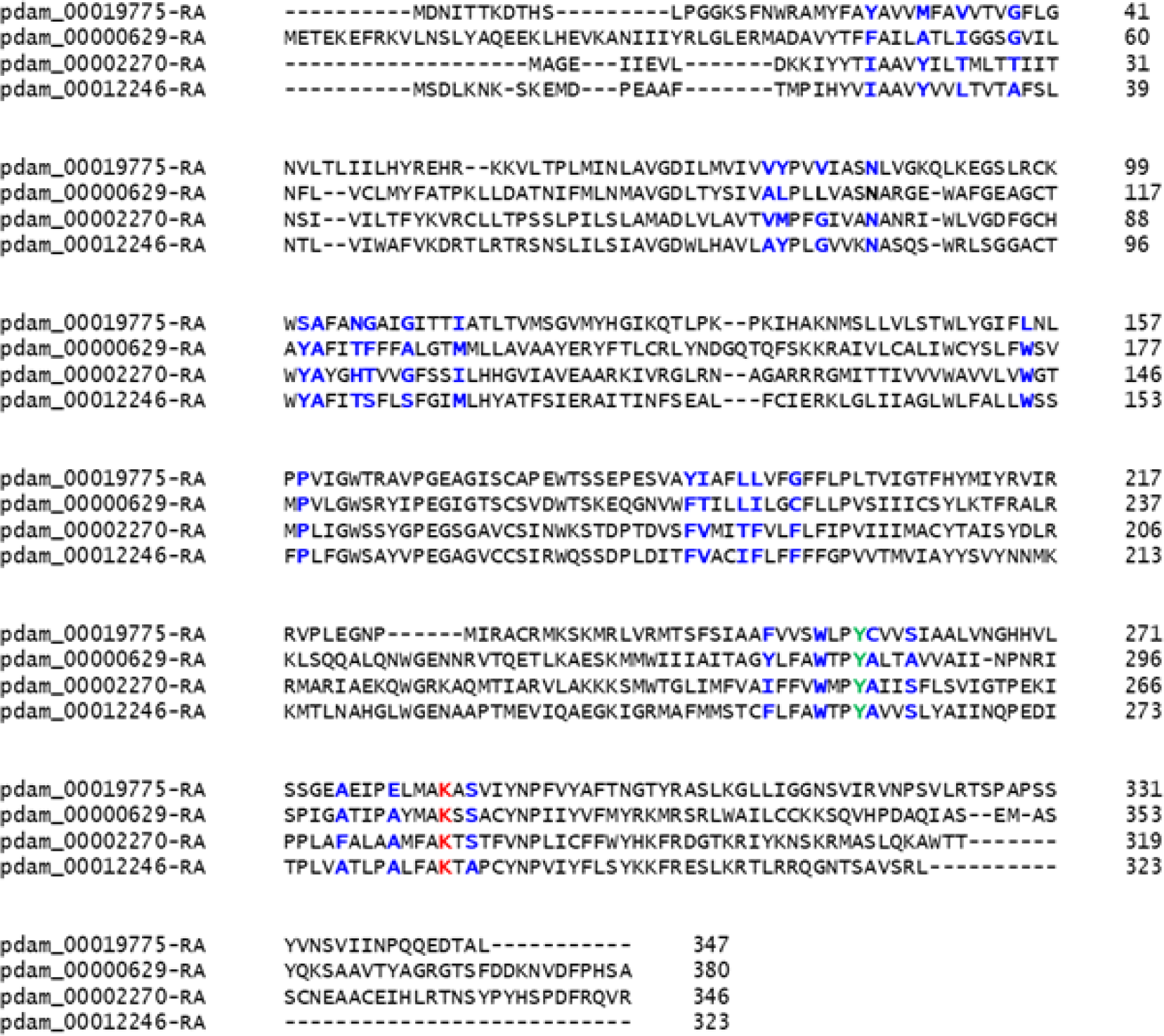
Clustal omega multiple sequence alignment of opsin homologs.Residues involved in active site formation are highlighted in blue.

**Supplementary Table S1.**
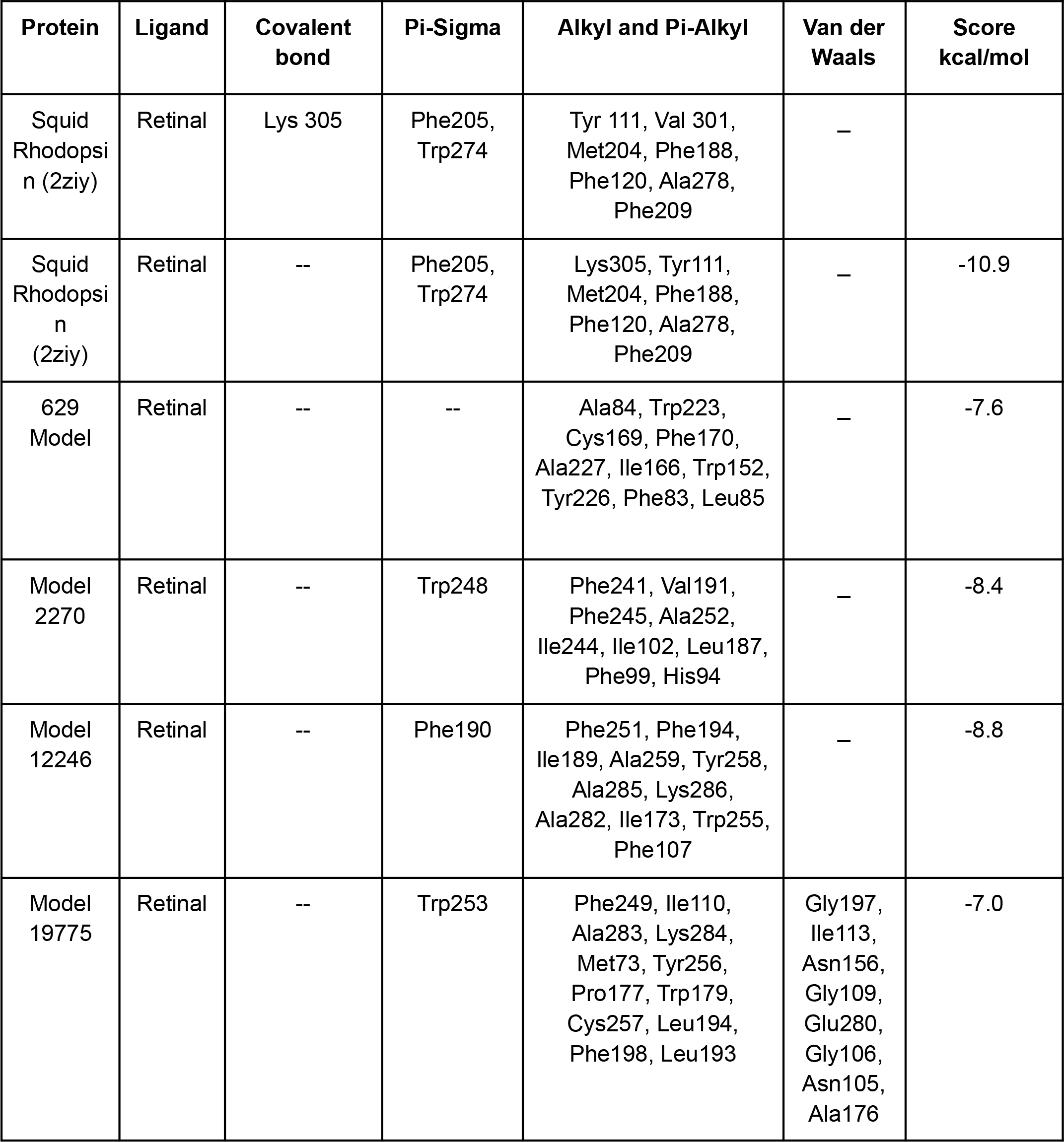
The details of residues and their molecular interactions with the retinal active site in squid rhodopsin and coral putative rhodopsin proteins.

| Protein | Ligand | Covalent bond | Pi-Sigma | Alkyl and Pi-Alkyl | Van der Waals | Score kcal/mol |
| --- | --- | --- | --- | --- | --- | --- |
| Squid Rhodopsin (2ziy) | Retinal | Lys 305 | Phe205, Trp274 | Tyr 111, Val 301, Met204, Phe188, Phe120, Ala278, Phe209 | – |  |
| Squid Rhodopsin (2ziy) | Retinal | -- | Phe205, Trp274 | Lys305, Tyr111, Met204, Phe188, Phe120, Ala278, Phe209 | – | -10.9 |
| 629 Model | Retinal | -- | -- | Ala84, Trp223, Cys169, Phe170, Ala227, Ile166, Trp152, Tyr226, Phe83, Leu85 | – | -7.6 |
| Model 2270 | Retinal | -- | Trp248 | Phe241, Val191, Phe245, Ala252, Ile244, Ile102, Leu187, Phe99, His94 | – | -8.4 |
| Model 12246 | Retinal | -- | Phe190 | Phe251, Phe194, Ile189, Ala259, Tyr258, Ala285, Lys286, Ala282, Ile173, Trp255, Phe107 | – | -8.8 |
| Model 19775 | Retinal | -- | Trp253 | Phe249, Ile110, Ala283, Lys284, Met73, Tyr256, Pro177, Trp179, Cys257, Leu194, Phe198, Leu193 | Gly197, Ile113, Asn156, Gly109, Glu280, Gly106, Asn105, Ala176 | -7.0 |

